# Human neuromodulatory assembloids to study serotonin signaling and disease

**DOI:** 10.64898/2026.03.08.710407

**Authors:** Sabina Kanton, Xiangling Meng, Chunyang Dong, Fikri Birey, Dong Wang, Noah Reis, Se-Jin Yoon, Ji-Il Kim, James P. McQueen, Noriaki Sakai, Seiji Nishino, John R. Huguenard, Sergiu P. Paşca

**Affiliations:** Department of Psychiatry and Behavioral Sciences, Stanford University, Stanford, CA, USA; Stanford Brain Organogenesis, Wu Tsai Neurosciences Institute and Bio-X, Stanford University, Stanford, CA, USA; Department of Neurology and Neurological Sciences, Stanford University, Stanford, CA 94305, USA

## Abstract

Neuromodulators influence critical functions of the developing human brain and regulate behavioral states. Dysfunction of neuromodulatory systems is often involved in neuropsychiatric disease and many drugs for these conditions act on these signaling pathways. Recent advances in stem cell biology have made it possible to derive a wide range of cells across the developing human nervous system in regionalized organoids and to functionally integrate them into assembloids, however they currently do not systematically incorporate neuromodulation. Here, we generated human midbrain-hindbrain organoids (hMHO) from human induced pluripotent stem (hiPS) cells and fused them with human cortical organoids (hCO) to form neuromodulatory assembloids (hNMA). We focus on serotonin (5-hydroxytryptamine, 5-HT) as one key neuromodulator and found characteristic gene expression patterns and electrophysiological properties of serotonergic neurons (5-HT neurons) in the hMHO. In hNMA, 5-HT neurons projected into hCO, released 5-HT and modulated cortical network activity. To explore the applicability of this system in human disease, we studied 22q11.2 deletion syndrome (22q11.2DS), a common microdeletion associated with high risk for neuropsychiatric disease and defects in 5-HT signaling. We found aberrant 5-HT dynamics in hNMA from patient hiPS cell lines that were rescued by administration of a selective serotonin reuptake inhibitor (SSRI). Taken together, hNMA can be used to study human 5-HT dynamics and uncover disease phenotypes which could facilitate therapeutic development.

## Introduction

Neuromodulators including serotonin (5-HT), dopamine and norepinephrine, are involved in a broad spectrum of neuropsychiatric disorders such as anxiety, depression and schizophrenia and are among the main neurotransmitter systems targeted by drugs used in neuropsychiatry. Serotonin is one of the earliest neurotransmitters to emerge during central nervous system (CNS) development^1^. Starting from gestational week 5 (GW5), 5-HT is almost exclusively generated by serotonergic neurons (5-HT neurons) in the raphe nuclei located in the midbrain-hindbrain region^2,3^. These neurons send projections to almost all regions of the CNS to modulate neural activity and are critical in early cortical network formation^4,5^. The 14 subtypes of 5-HT receptors are prime targets for drug development for neuropsychiatric diseases^6^ and are among the top divergently expressed genes between human brain and mouse brain^7^ highlighting the necessity to study serotonergic neuromodulation in a human genetic context.

Stem cell-derived models of human brain development such as organoids recapitulate a wide range of neuronal diversity in the developing CNS^8^, and integrating regionalized organoids into assembloids allows for the study of functional connectivity via projections and cell migration in complex multi-region interactions^9,10^. Although assembloids allow to study signaling inputs and outputs, controlled neuromodulation of neural networks in assembloids is currently lacking.

Here, we introduce a human organoid model of the midbrain-hindbrain region (hMHO) which contains neuromodulators such as 5-HT and dopamine and which transcriptionally mapped to midbrain-hindbrain identities. We focus on 5-HT as a neuromodulator and used hMHO to establish a human neuromodulatory assembloid (hNMA) via integration with human cortical organoids (hCO). The serotonergic neurons in the hMHO produce and secrete 5-HT and anatomically and functionally connected to hCO which we demonstrated via 5-HT sensor imaging and optogenetic manipulation in conjunction with electrophysiology. Moreover, long-term fusion of hMHO to hCO in assembloids led to changes in network activity in hCO.

Lastly, we applied the hNMA platform to study 22q11.2 deletion syndrome (22q11.2DS) which is the most common microdeletion in humans and is one of the highest genetic risk factors for neuropsychiatric conditions such as autism spectrum disorder (ASD) and psychosis^11,12^. We used the hNMA platform and a genetically encoded 5-HT sensor to study 22q11.2DS and discovered disease phenotypes in hCO in the patient-derived hNMA but not in hMHO or hCO alone that were effectively modulated by an SSRI. This highlights the potential for the broad applicability of this system to study disease mechanisms in the context of neuromodulation and to develop new interventions for neuropsychiatric disorders.

## Results

### Generation and characterization of human midbrain-hind-brain (hMHO) organoids from hiPS cells

Specification of the midbrain-hindbrain region relies on patterning of Sonic Hedgehog (SHH) and WNT pathways in addition to fibroblast growth factors (FGFs) such as FGF4 and FGF8 located at the midbrain-hindbrain boundary (Figure 1A)^2,13^. Previous protocols to generate 5-HT neurons in 2D cultures from hiPS cells and caudal hindbrain neurons have been described^14,15^. Here, we modulated SHH pathways using Smoothened agonist (SAG) and WNT pathways using CHIR to obtain patterning along the rostral-caudal and dorso-ventral axes in addition to adding FGF4 to specify the midbrain-hindbrain boundary region (Figure 1B). To validate patterning of the regional identities, we verified expression of the early hindbrain markers *GATA2* and *GATA3* at 19-22 days and absence of forebrain marker *FOXG1* at day 50-60 using RT-qPCR (Figure S1).

**Figure 1:**
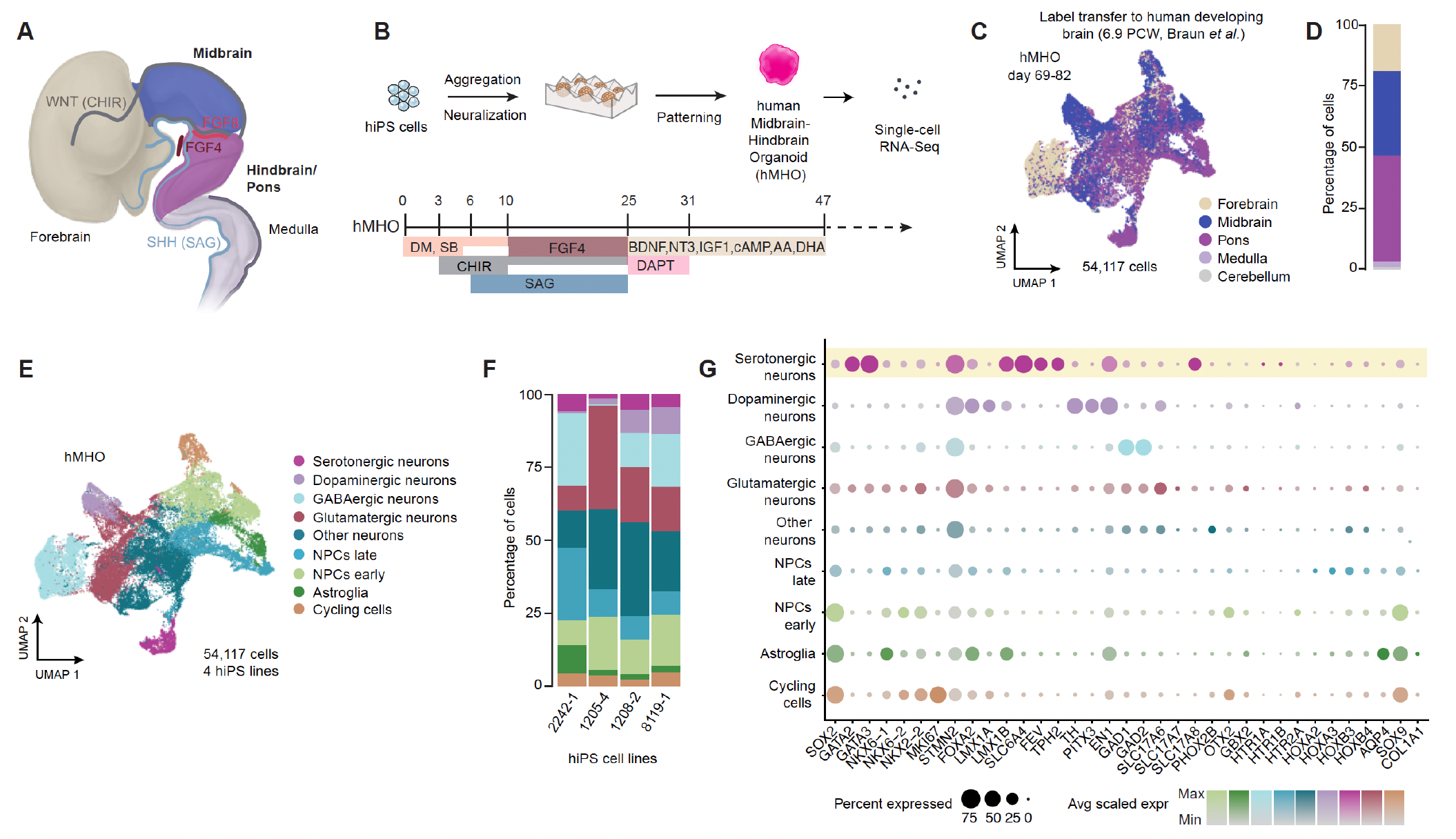
Generation and characterization of human midbrain-hindbrain organoids (hMHO). **(A)** Schematic of the developing brain and small molecules used to pattern midbrain-hindbrain regions (adapted from Wurst and Bally-Cuif^13^). **(B)** Schematic and recipe for generating hMHO. **(C)** Single-cell RNA-seq analysis of 54,117 cells (4 hiPS cell lines, 4 differentiation experiments, day 69-82) in hMHO and label transfer of cells to brain regions of PCW 6.9 human developing brain tissue. **(D)** Proportions of cells in hMHO based on label transfer to PCW 6.9 human developing brain tissue. **(E)** Cell type annotation in hMHO. NPCs: neural progenitor cells. **(F)** Proportions of cell types in hMHO across 4 different hiPS cell lines. **(G)** Dotplot of marker gene expression in hMHO across different cell types. The size of the circles represents the percentage of cells expressing the marker gene and the color represents scaled gene expression.

We used single-cell RNA-sequencing (scRNA-seq) to profile the cellular diversity in hMHO and sequenced 54,117 cells across 4 hiPS cell lines between day 69 and 82 of differentiation. To verify their regional identity, we mapped the cells to primary human developing brain transcriptomic data at PCW 6.9^16^ (Figure 1C) using Seurat label transfer^15^ and observed that more than 75% of cells overlap with midbrain and hindbrain (pons) regional identities (Figure 1C). We found distinct cell type clusters of hindbrain progenitor cells (*SOX2*^*+*^, *GATA2*^*+*,^ *GATA3*^*+*^, *NKX6-1*^*+*^, *NKX2-2*) and neuronal cell types including early neurons (*STMN2*^*+*^) and more mature neurons with various neurotransmitter or neuromodulator signatures such as glutamatergic (*SCL17A6*^*+*^, *SCL17A7*^*+*^, *SLC17A8*^*+*^), GABAergic (*GAD1*^*+*^, *GAD2*^*+*^), dopaminergic (*TH+*) and serotonergic markers (*TPH2+, SLC6A4*^*+*^, *FEV*^*+*^), as well as cycling progenitor cells (*MKI67*^*+*^) and astroglia (*SOX9*^*+*^, *AQP4*^*+*^) (Figure 1E, G, Figure S2A). The proportions of cell types were overall consistent across four different hiPS cell lines (Figure 1F).

### Characterization of serotonergic neurons in hMHO

The hMHO protocol produced a group of 5-HT neurons that expressed the canonical markers *TPH2* (tryptophan hydroxylase 2), which encodes the key enzyme for synthesis of 5-HT in the brain, *SLC6A4* (5-HT transporter), encoding the serotonin transporter SERT, and *FEV*, which encodes a transcription factor that is essential for 5-HT neuron formation^17^ (Figure 1G, 2A, Figure S1, S2). We mapped these neurons to the primary human developing brain transcriptome reference at 6.9 PCW^16^ using Seurat label transfer^18^ and observed that more than 85% of all the cells mapped to midbrain/hindbrain (pons) regional identities (Figure 2B). We also identified previously described subtype marker expression patterns^19^ (Figure S2B-D) for 5-HT neurons. When we mapped the cells to a mouse E13 spatial reference using VoxHunt^20^, we detected overlap with the midbrain/hindbrain region that furthermore aligned with the spatial expression pattern of *Tph2* (Figure 2C). We performed immunohistochemical (IHC) staining of TPH2 in hMHO cryosections to test for the presence of this marker at the protein level and observed TPH2^+^ cells throughout the hMHO (Figure 2D, Figure S3A, B). We checked for the presence of 5-HT using IHC in hMHO cryosections and observed 5-HT^+^ cells (Figure 2E, Figure S3C). Moreover, we used high performance liquid chromatography (HPLC) to measure the levels of 5-HT in hMHO and hCO and detected 5-HT in hMHO but not in hCO (Figure 2F).

**Figure 2:**
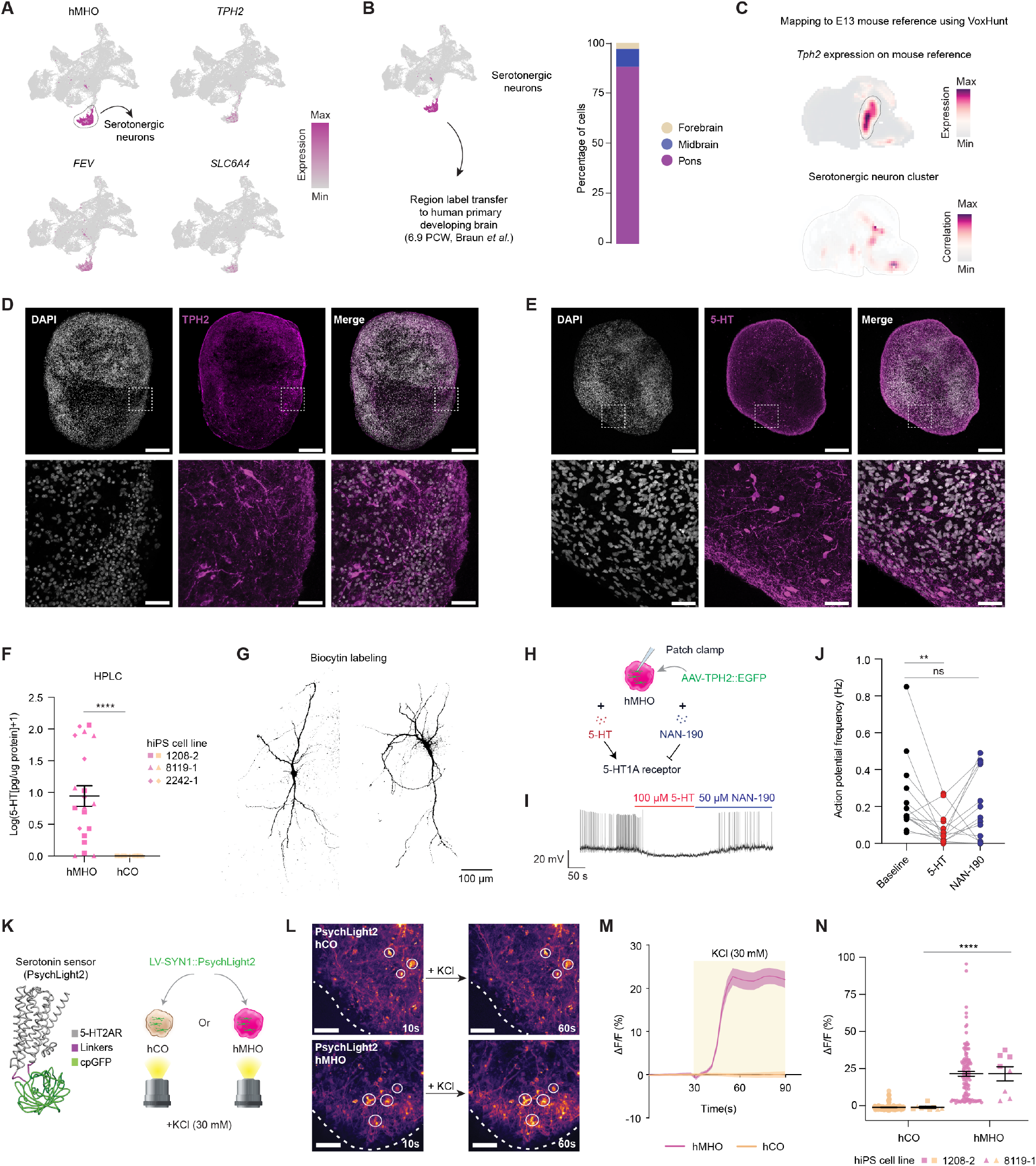
Characterization of 5-HT neurons in hMHO. **(A)** Canonical markers of 5-HT neurons in single-cell RNA-seq data of hMHO. **(B)** Proportions of brain regions mapping to 5-HT neurons based on label transfer of PCW 6.9 primary developing human brain. **(C)** *Tph2* expression in mouse E13 spatial atlas and projection of 5-HT neurons onto mouse reference using VoxHunt^20^. **(D)** Immunohistochemical staining of TPH2 in hMHO (day 84, scale bar = 250 *µ*m (top) and 40 *µ*m (bottom)). **(E)** Immunohistochemical staining of 5-HT in hMHO (day 83). **(F)** HPLC measurements of 5-HT in hMHO and hCO (hMHO: 20 samples, 13 differentiation experiments, hCO: 9 samples, 4 differentiation experiments; 3 hiPS cell lines, day 91-183. Mann-Whitney test, ****p<0.0001). **(G)** Biocytin labeling of AAV-TPH2::EGFP labeled 5-HT neurons (day 157, one iPS cell line, one differentiation experiment). **(H)** Schematic of patch clamp recording of 5-HT neurons (TPH2::EGFP^+^ cells) in hMHO at baseline and during addition of 5-HT and NAN-190. **(I)** Example trace of action potentials during addition of 5-HT (100 *µ*m) and NAN-190 (50 *µ*m, 5-HT1A receptor blocker). **(J)** Action potential frequencies of TPH2::EGFP^+^ neurons during baseline and addition of 5-HT and NAN-190 (14 cells, 1 hiPS cell line, 3 differentiation experiments. Friedman test *p = 0.0124, with Dunn’s multiple comparison testing: baseline vs. 5-HT **p = 0.0092, baseline vs. NAN-190 p = 0.7902, ns: non-significant). **(K)** Structure of serotonin sensor PsychLight2 (adapted from Dong *et al*^26^ with permission from Elsevier publishing, License #: 6152850897994) and schematic of serotonin sensor imaging. **(L)** Example images of PsychLight2 imaging in hCO and hMHO during KCl stimulation at baseline (10s) and during KCl stimulation (60s) (day 172, scale bar = 200 *µ*m). **(M)** Quantification of temporal dynamics of serotonin sensor fluorescence changes in hMHO and hCO (hCO: 9 organoids, hMHO: 8 organoids, 2 hiPS cell lines, 3 differentiation experiments). **(N)** Change in fluorescence intensity for PsychLight2 in hCO and hMHO after 30 s of KCl stimulation across different iPS cell lines (hCO: n = 135 ROIs (left), 9 organoids (right), hMHO: 120 ROIs (left), 8 organoids (right), 2 hiPS cell lines, 3 differentiation experiments for hCO and hMHO. Mann-Whitney test, ****p<0.0001).

To visualize 5-HT neurons in live *in vitro* hMHO cultures, we designed an adeno-associated virus (AAV) expressing EGFP under the control of a *TPH2* promoter based on a previously described promoter sequence^21^ and observed overlap between TPH2 and EGFP protein signal using IHC on hMHO cryosections (Figure S3D). We then filled EGFP^+^ neurons in hMHO using biocytin and visualized them via streptavidin conjugated to Dylight549 and observed that they exhibited complex multipolar morphologies with long processes extending over several hundred microns (Figure 2G). A characteristic property of 5-HT neurons is their reduced firing in the presence of 5-HT which is mediated via the somatodendritic 5-HT1A autoreceptor^22,23^. We tested this by performing patch clamp electrophysiology on EGFP^+^ cells in hMHO while perfusing with 5-HT and observed a significant decrease in action potential frequency (Figure 2H-J). This effect was partially rescued by perfusing with NAN-190, a 5-HT1A receptor blocker, thereby (Figure 2J) indicating autoinhibition properties in 5-HT neurons in hMHO.

Serotonergic neurons have been described to exhibit characteristic electrophysiological properties compared to non-serotonergic neurons such as increased action potential (AP) half width^24,25^. To investigate this, we performed patch clamp on TPH2::EGFP^+^ cells in hMHO and compared them to SYN1::mCherry^+^ (Synapsin-1) cortical neurons in hCO (Figure S4). We indeed detected a significant difference in action potential half width, with TPH2::EGFP^+^ cells exhibiting an average duration of 3.42 ± 0.15 ms compared to 2.59 ± 0.13 ms in SYN1::mCherry^+^ cells (Figure S4E). We also observed a significant difference in AP threshold between serotonergic and cortical neurons, but no difference in peak AP amplitude, resting membrane potential or capacitance (Figure S4F-I).

To monitor the presence of 5-HT in live hMHO at high temporal and spatial resolution using confocal imaging, we implemented a genetically encoded 5-HT sensor under the control of the SYN1 promoter (PsychLight2)^26^ and infected the construct via lentiviral (LV) transduction (Figure 2K). We first validated 5-HT sensor responses in hCO that do not produce 5-HT endogenously by perfusing with a range of 5-HT concentrations. We observed a wide dynamic range of detection across four orders of magnitude and up to 100 nM of 5-HT (Figure S5). Furthermore, the fluorescence signal was significantly reduced to baseline levels using a 5-HT2A receptor antagonist (EMD281014) that competes with 5-HT for binding to the sensor^26^. We then transduced hCO and hMHO with LV-SYN1::PsychLight2 and measured 5-HT sensor signal when stimulating with KCl (Figure 2K). We detected significant fluorescence signal increases in hMHO but not in hCO (Figure 2L) and the signal lasted over an extended period (Figure 2M) which was consistent across organoids generated from multiple hiPS cell lines (Figure 2N). Taken together, these results suggest that hMHO contains functional 5-HT neurons that secrete 5-HT indicating that this platform can be used to study human serotonergic neuromodulation.

### Building a human neuromodulatory assembloid (hNMA) with functional connectivity

Serotonergic neurons project into the forebrain and modulate the development and activity of cortical networks^5^. To model the influence of 5-HT on cortical networks, we generated an assembloid that we named human neuromodulatory assembloid (hNMA) by fusing hMHO to hCO around day 60 of differentiation. To visualize projections between the organoids, we transduced hCO with AAV-SYN1::mCherry and hMHO using the AAV-TPH2::EGFP prior to assembly (Figure 3A). We then monitored projections using confocal live imaging after 2, 4 and 8 weeks of fusion by quantifying the area of EGFP^+^ signal in hCO and mCherry^+^ signal in hMHO. We detected bilateral projections between hMHO and hCO that increased over two months of fusion, confirming anatomical connection within the hNMA (Figure 3B-D). To test whether 5-HT is released from hMHO into hCO in hNMA, we transduced hCO with the 5-HT sensor (LV-SYN1::PsychLight2) and fused them to hMHO around day 60 (Figure 3E). We then investigated 5-HT release into hCO in hNMA using KCl stimulation and detected an immediate and lasting increase in fluorescence intensity in hCO, indicating 5-HT release from hMHO (Figure 3F, G). To test whether organoids need to be functionally integrated into hNMA for measurable 5-HT release, we placed unfused hCO and hMHO at a distance within a microscopy field and stimulated with KCl. We did not detect an increase in fluorescence signal in unassembled hCO compared to hCO in hNMA (Figure 3F-H) indicating that anatomical connection between the organoids in an assembloid is necessary to obtain measurable 5-HT levels in hCO.

**Figure 3:**
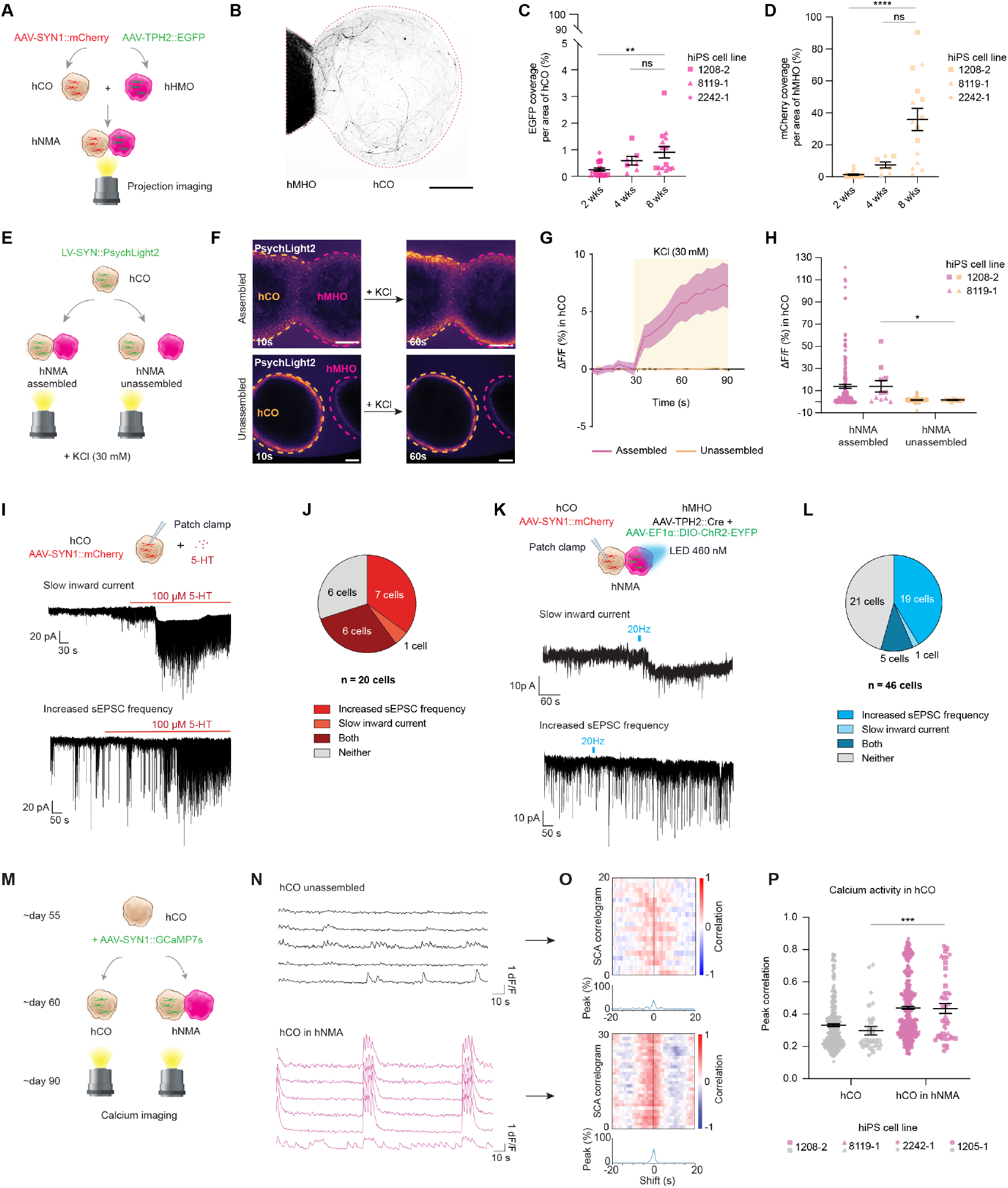
Anatomical and functional connectivity in hNMA. **(A)** Schematic of imaging of bilateral projections in hNMA. **(B)** Example image of TPH2::EGFP projections from hMHO to hCO in hNMA after ~4 weeks of fusion (scale bar = 500 *µ*m). **(C)** Quantification of TPH2-EGFP^+^ projections into hCO (n = 41 assembloids, 3 hiPS cell lines, 4 differentiation experiments. Kruskal-Wallis test, p** = 0.0033 with Dunn’s multiple comparison testing: 2 wks vs. 8 wks **p = 0.0025, 4 wks vs. 8 wks p > 0.9999). **(D)** Quantification of SYN1-mCherry^+^ projections into hMHO (n = 41 assembloids, 3 hiPS cell lines, 4 differentiation experiments. Kruskal-Wallis test, p**** < 0.0001 with Dunn’s multiple comparison testing: 2 wks vs. 8 wks ****p < 0.0001, 4 wks vs. 8 wks p = 0.2911). **(E)** Schematic of serotonin sensor imaging in hNMA and unassembled hCO and hMHO. **(F)** Example images of serotonin sensor imaging of fused and unfused hNMA at base line (10s) and after KCl stimulation (60s) (scale bar = 200 *µ*m). **(G)** Temporal dynamics of serotonin sensor fluorescence signal in unassembled hCO and hCO in hNMA (assembled: n = 11 assembloids, unassembled n = 10 pairs of organoids, 2 hiPS cell lines, 3 differentiation experiments). **(H)** Quantification of serotonin sensor signal at maximum fluorescence intensity across multiple hiPS cell lines (assembled: n = 165 ROIs (left), 11 assembloids (right); unassembled: 150 ROIs (left), 10 pairs of organoids (right); 2 hiPS cell lines, 3 differentiation experiments. Mann-Whitney test, *p=0.0159). **(I)** Example patch clamp traces and response types of SYN1::mCherry^+^ cells in hCO during 5-HT addition. **(J)** Quantification of response types in hCO during 5-HT administration (20 cells, 2 hiPS cell lines, 3 differentiation experiments, 157-180d). **(K)** Example patch clamp traces and response types in SYN1::mCherry^+^ cells in hCO in hNMA during optogenetic stimulation of hMHO. **(L)** Quantification of response types in hCO during optogenetic stimulation in hNMA (46 cells, one hiPS cell line, 3 differentiation experiments, 142-200d). **(M)** Schematic of calcium imaging in hCO and hNMA. **(N)** Example calcium imaging traces in unassembled hCO and hCO in hNMA. **(O)** SCA correlograms of unassembled hCO and hCO in hNMA. **(P)** Peak correlations from SCA analysis in hCO and hCO in hNMA in multiple hiPS cell lines (hCO: 318 cells (left), 30 organoids (right); hCO in hNMA: 463 cells (left), 44 organoids (right), 3 hiPS cell lines, 3 differentiation experiments. Mann-Whitney-U test ***p = 0.0004).

Serotonin modulates the activity of cortical neurons acutely by binding to a variety of 5-HT receptors^5^. To investigate the dynamic changes in cortical neurons upon receiving 5-HT, we first tested the effect of 5-HT perfusion on hCO labeled with AAV-SYN1::mCherry using patch clamp electrophysiology (Figure 3I). By recording from mCherry^+^ cells, we observed different types of previously reported responses^27^, such as an increase in sEPSC (spontaneous excitatory postsynaptic current) frequency, a slow inward current response, or a combination of both responses (Figure 2I, J). We then transduced hMHO with Channelrhodopsin in TPH2^+^ cells (AAV-TPH2::Cre + AAV-EF1α::DIO-ChR2-EYFP) and fused them to hCO that were transduced with SYN1::mCherry to stimulate 5-HT release using optogenetics. We performed patch clamp recordings in SYN1::mCherry^+^ neurons while optogenetically stimulating hMHO using a train stimulation protocol at 20 Hz^28^ (Figure 3K). We detected similar types of responses in hCO during optogenetic stimulation in hNMA as in hCO with 5-HT perfusion when comparing to baseline activity such as sEPSC frequency increases and inward currents (Figure 3K, L). Overall, the responses were similar when perfusing 5-HT in hCO or during optogenetic activation of hMHO in hNMA, however the proportion of non-responding cells was higher during optogenetic activation, and we observed a higher proportion of increased sEPSC compared to inward currents in hNMA (Figure 3J, 3L). Overall, these findings suggest intact anatomical and functional connectivity in hNMA and recapitulate aspects of neuromodulation in cortical neurons.

We next investigated the effect of long-term fusion in hNMA on cortical neuron activity to study potential chronic effects of 5-HT. By transducing hCO with AAV-SYN1::GCaMP7s to express a genetically encoded calcium sensor to visualize calcium transients, we compared cortical neuron activity in unfused hCO with activity in hCO fused to hMHO after 1 month of integration using confocal live imaging (Figure 3M). We quantified the calcium transients and observed more correlated signals in hCO fused to hMHO compared with unassembled hCO (Figure 3N). We further scrutinized calcium activity using scaled correlation analysis (SCA) (Figure 3O) which allowed for an improved quantification of synchronous activity in neural circuits by reducing the contribution of low-frequency noise signals^29,30^. This analysis revealed a significant increase in correlated activity between cells in hCO integrated with hMHO compared to hCO alone (Figure 3P) suggesting fusion-induced changes in neural activity and increased synchrony. These observations imply long-term effects of neuromodulation on cortical network activity that can be modeled in hNMA.

### Serotonin dynamics in hNMA derived from patients with 22q11.2 deletion syndrome

Finally, we investigated the potential effects of 22q11.2DS on 5-HT neuromodulation dynamics in hNMA since this deletion has been associated with reduced 5-HT levels in affected individuals^31^ along with symptoms suggesting 5-HT dysregulation^11,12^. To address this question, we used five 22q11.2DS patient-derived cell lines^32^ (Figure 4A) and confirmed deletion in the 22q11.2 region using SNP arrays (Figure 4B). We generated hNMA from control and 22q11.2DS cell lines and investigated 5-HT dynamics using 5-HT sensor imaging in the hCO side of hNMA in conjunction with chemical depolarization via KCl (Figure 4C). We observed a significant difference between control and 22q11.2DS hNMA, with controls exhibiting on average a 9.8% increase in fluorescence intensity compared to 3.3% in 22q11.2DS, suggesting lower levels of 5-HT in 22q11.2DS at maximum signal intensity (Figure 4C). To dissect the contribution of the different regions to the observed phenotype, we conducted the same experiment in hCO and hMHO separately. We did not observe any significant differences between control and 22q11.2DS in either of the regions (Figure 4D, E). We tested whether defects in anatomical projection between hMHO and hCO might contribute to the observed phenotype, hypothesizing that differences in the number of projections could lead to different levels of 5-HT in hCO in 22q11.2DS. To address this, we performed projection imaging on the hCO side in hNMA around 1 month after assembly with hMHO transduced with AAV-TPH2::EGFP by measuring the percentage of EGFP coverage per area in hCO (Figure 4F). Interestingly, we did not observe any significant differences in the area covered by projections in 22q11.2DS compared to control, suggesting no major defects in anatomical connectivity in hNMA derived from 22q11.2DS patient cell lines (Figure 4G).

**Figure 4:**
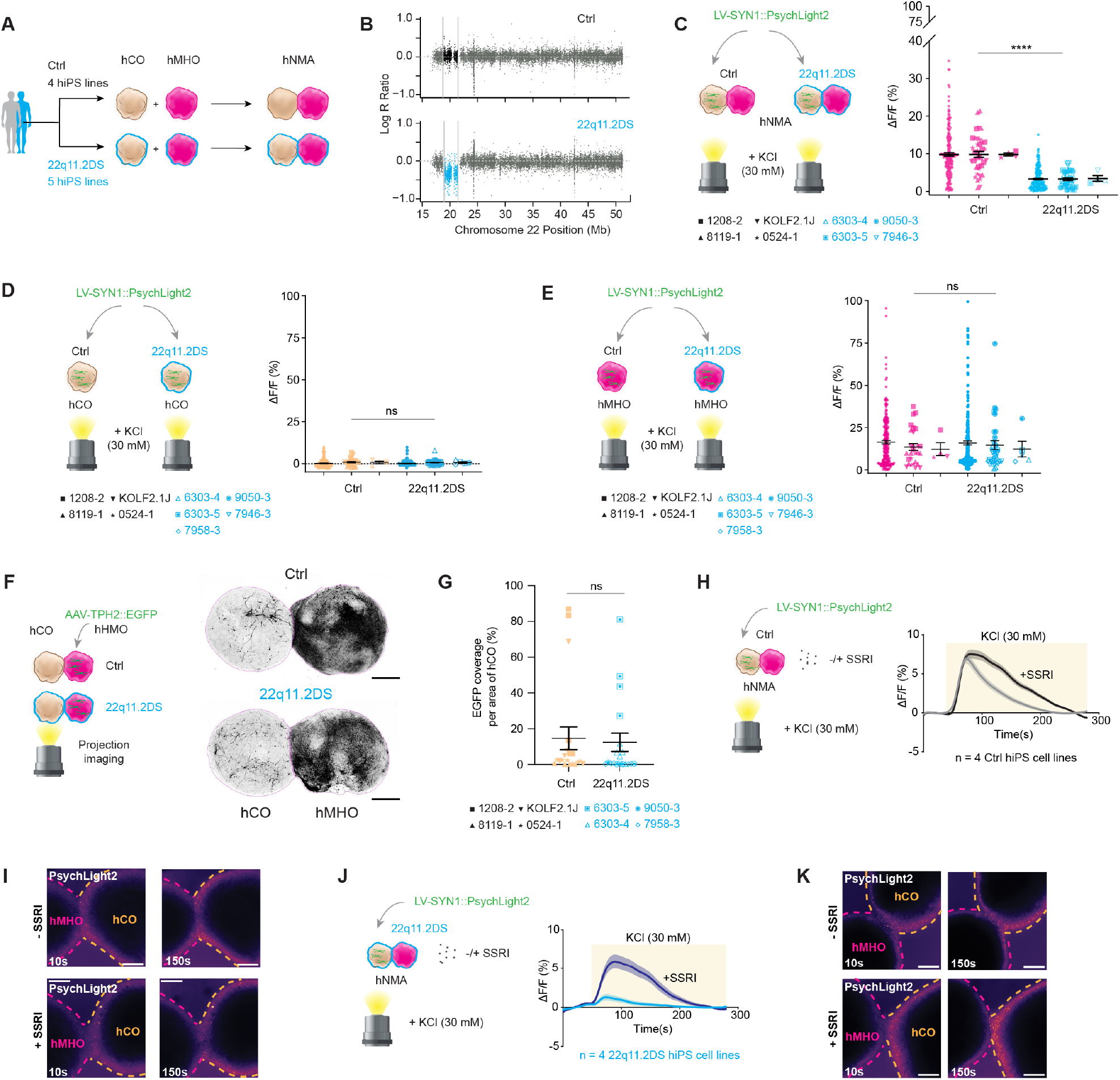
Disease modeling of 22q11.2DS using hMHO and hNMA. **(A)** Patient-derived 22q11.2DS and control hiPS cell lines for generating hCO, hMHO and hNMA. **(B)** Example SNP array of one control and 22q11.2DS hiPS cell line (22q11.2DS: 6303-5, Ctrl: 1208-2). **(C)** PsychLight2 imaging in hCO from 22q11.2DS and control hNMA during KCl stimulation (Ctrl: 200 ROIs (left), 40 assembloids (middle), 4 hiPS cell lines (right), 3 differentiation experiments; 22q11.2DS: 175 ROIs (left), 35 assembloids (middle), 4 hiPS cell lines (right), 3 differentiation experiments. Unpaired *t*-test with Welch’s test, ****p<0.0001). **(D)** PsychLight2 imaging in 22q11.2DS and control hCO during KCl stimulation (Ctrl: 225 ROIs (left), 27 organoids (middle), 4 hiPS cell lines (right), 3 differentiation experiments; 22q11.2DS: 300 ROIs (left), 36 organoids (middle), 5 hiPS cell lines (right), 3 differentiation experiments. Unpaired t-test with Welch’s test, p = 0.9690). **(E)** PsychLight2 imaging in 22q11.2DS and control hMHO during KCl stimulation (Ctrl: 220 ROIs (left), 28 assembloids (middle), 4 hiPS cell lines (right), 3 differentiation experiments; 22q11.2DS: 280 ROIs (left), 32 assembloids (middle), 5 hiPS cell lines (right), 3 differentiation experiments. Unpaired t-test with Welch’s test, p = 0.7334). **(F)** Imaging of TPH2::EGFP^+^ projections from hMHO to hCO in hMNA and examples of TPH2::EGFP^+^ signals in control and 22q11.2DS hNMA (scale bar = 500 *µ*m). **(G)** Quantification of TPH2::EGFP^+^ projections from hMHO to hCO in hMNA after 30-40 days of fusion (Ctrl: 20 assembloids, 2 differentiation experiments, 4 hiPS cell lines; 22q11.2DS: 19 assembloids, 2 differentiation experiments, 4 hiPS cell lines. Mann-Whitney test, p = 0.4955). **(H)** PsychLight2 imaging in hCO from control hNMA with SSRI treatment during KCl stimulation (without SSRI: 18 assembloids, 4 hiPS cell lines, 3 differentiation experiments; with SSRI: 20 assembloids, 4 hiPS cell lines, 3 differentiation experiments). **(I)** Representative images of PsychLight2 imaging in control hNMA with and without SSRI addition. **(J)** PsychLight2 imaging in hCO 22q11.2DS hNMA with SSRI treatment during KCl stimulation (without SSRI: 19 assembloids, 4 hiPS cell lines, 3 differentiation experiments; with SSRI: 19 assembloids, 4 hiPS cell lines, 3 differentiation experiments). **(K)** Representative images of PsychLight2 imaging in 22q11.2DS hNMA with and without SSRI addition.

Selective serotonin reuptake inhibitors (SSRIs) are one of the main pharmacological interventions to treat neuropsychiatric symptoms in 22q11.2DS^33^ and we speculated that they might rescue the reduced 5-HT levels in hNMA derived from 22q11.2DS patients. To this end, we performed 5-HT sensor imaging on the hCO side of control and 22q11.2DS hNMA during KCl stimulation with and without the addition of fluoxetine, a commonly used SSRI. Fluoxetine administration in control hNMA did not change the maximum intensity of fluorescence but led to a prolonged fluorescence signal in hCO (Figure 4H, I). Interestingly, in the 22q11.2DS condition, SSRI addition increased the amplitude and the duration of the 5-HT induced fluorescence in hCO (Figure 4J, K). When quantifying the cumulative fluorescence signal as area under the curve (AUC), the addition of the SSRI led to a partial rescue of the observed phenotype in 22q11.2DS by increasing the cumulative 5-HT signal to levels similar to controls (Figure S6). These results underscore the feasibility of modeling 22q11.2DS neuromodulation defects in a human assembloid model and the applicability of this model system to study aberrant neuromodulation in disease states.

## Discussion

Many drugs used for neuropsychiatric conditions target neuromodulator systems such as 5-HT, dopamine and norepinephrine. Although stem cell-based models have been able to recapitulate long-range projections and migration in assembloids^9,10^ neuromodulation via 5-HT has not been specifically addressed. Here, we generated an organoid of midbrain-hindbrain identity containing different neurotransmitters and neuromodulators such as 5-HT and dopamine. We recapitulated characteristic properties of 5-HT neurons such as 5-HT production as measured by HPLC and a genetically encoded 5-HT sensor (PsychLight2), and electrophysiological properties using patch clamp measurements. Generation of hNMA by fusing hMHO with hCO resulted in long range serotonergic projections and functional connectivity via 5-HT release as visualized by 5-HT sensor imaging and patch clamp recording with optogenetic stimulation. Long-term fusion of these hNMA revealed synchronized cortical network activity. Delineating how this effect might be mediated by 5-HT, for example, via 5-HT receptors or epigenetically induced modifications such as serotonylation of histones^34^, could identify the underlying mechanism of the observed network changes.

The current model has several limitations that should be addressed to further improve this hNMA system. Serotonin is known to modulate the function of GABAergic neurons^35^ which are largely absent in hCO. These could be integrated by fusing subpallial organoids^9^ to hCO to generate a 3-part assembloid to more faithfully recapitulate cortical circuits that incorporate inhibitory and excitatory neurons. The hMHO differentiation protocol could be expanded more caudally to incorporate 5-HT neurons that project to the hindbrain and spinal cord in order to explore projection specificity of 5-HT neurons in various assembloid configurations^15^.

We applied our model to study 22q11.2DS, one of the most common microdeletions in humans that is often associated with symptoms suggesting neuromodulator dysfunction such as psychosis^12^. 22q11.2DS has been previously studied in hCO and cortico-thalamic assembloids revealing hyperexcitability in cortical neurons^32^ and overgrowth of thalamic axons into the cortex^36^, respectively. However, the understanding of 5-HT in 22q11.2DS is currently limited despite indications of lower levels of 5-HT in individuals with 22q11.2DS^31^ and positive effects of early SSRI treatment to address psychiatric symptoms^33,37^. We detected phenotypes of 22q11.2DS by using 5-HT sensor imaging and found lower levels of 5-HT in hCO upon stimulation of hMHO, an effect that was rescued by a SSRI. The modular nature of the hNMA could help to further elucidate the mechanism of this phenotype by using mosaic assembloids of control and 22q11.2DS organoids. This would allow to dissect the contribution of hMHO-related differences in 5-HT release and potential hCO-related differences in 5-HT uptake via monoamine transporters^38^ or differences in 5-HT receptor signaling.

Going forward, the hNMA will enable investigation of the role of 5-HT in other brain regions, for example by integrating hMHO with striatal organoids^39^, or to study the effects of other neuromodulators such as dopamine on cortical networks. Neuromodulators that are not present in our model such as norepinephrine could be incorporated by modifying the current differentiation protocol. Taken together, this new platform holds promise for uncovering insights into neuromodulatory systems in the brain and could help establish and test new treatment opportunities at the cellular and molecular level.

## Supporting information

qPCR Primer Sequences

## Acknowledgements

We thank all members of the Pasca lab for helpful discussions. We thank M. Onesto for helpful discussions and assistance with projection imaging experiments, M.-Y. Li for guidance and useful discussions on patch clamp experiments, and Y. Miura for helpful discussions on assembloid experiments. We thank K. W. Kelley for providing processed data for the primary tissue data set from Braun *et al*. and M. V. Thete for help with qPCRs in the initial stages of the project. We thank the Stanford Gene Vector and Virus Core (RRID:SCR_023250) for providing AAVs and the Stanford Genomics Service Center for Bioanalyzer analysis for single-cell RNA-seq experiments.

This work was supported by an NIH grant (R01MH107800), the Stanford Big Idea Project on Brain Organogenesis (Wu Tsai Neurosciences Institute) (to S.P.P.), the Kwan Funds (to S.P.P.), the Senkut Funds (to S.P.P.), the Coates Foundation (to S.P.P.), the Ludwig Family Foundation (to S.P.P.), the Alfred E. Mann Foundation (to S.P.P.), an EMBO Postdoctoral Fellowship (ALTF 321-2021; to S.K.), a Stanford University Postdoctoral Fellowship (to X.M.), a Stanford Maternal & Child Health Research Institute (MCHRI) Postdoctoral Fellowship (to F.B.), and the American Epilepsy Society Postdoctoral Research Fellowship (to F.B.). S.P.P. is a CZ BioHub Investigator and a CZI Ben Barres Investigator.

## Author contributions

F.B., X.M., S.K., C.D., and S.P.P. conceived the study and designed experiments. F.B. and X.M. developed the hMHO differentiation protocol. X.M., N.R., S.K., S.Y., J.P.M, F.B., and C.D performed organoid culture. S.K. and F.B. performed single-cell RNA-seq experiments and library prep with help from X.M.. S.K. analyzed single-cell RNA-seq data with initial analysis by F.B.. N.R., J.P.M. and S.Y. performed RT-qPCR experiments with help from X.M., S.K., and F.B.. N.R. performed IHC with help from X.M., S.K., and J.P.M. D.W. and S.K. performed biocytin labeling experiments. X.M. designed and validated the AAV-TPH2::EGFP with help from N.R. and S.K.. S.K., C.D., and X.M. performed imaging of projections in assembloids, with help from N.R., and J.K. for analysis. D.W. performed patch clamp experiments. X.M. and N.R. performed calcium imaging experiments in assembloids. X.M. and N.R. analyzed calcium imaging data with help from J.K.. C.D. performed 5-HT sensor experiments and analyzed related data. N.S. and S.N. performed HPLC measurements. J.R.H. provided guidance on establishing and implementing functional assays. S.K., X.M., C.D., and S.P.P. prepared the manuscript with input from all the authors.

## Data availability

Raw and processed single-cell RNA-seq data for hMHO have been deposited to GEO (accession: GSE310824).

## Code availability

Python scripts for SCA^29,30^ analysis can be found at Zenodo: https://doi.org/10.5281/zenodo.13921409

## Materials and Methods

### hiPS cell culture

Human iPS cell lines (1208-2, 8119-1, 2242-1, 1205-4, 0524-1, 6303-4, 9050-3, 6303-5, 7946-3, 7958-3^32^ and KOLF2.1J^40^) were maintained in E8 in vitronectin coated cell culture plates. At around 80% confluency, cells were passaged every 4-5 days and were detached using 0.5 mM EDTA as previously described^41^. Cells were tested for the presence of mycoplasma using a PCR-based kit (abm, G238). SNP arrays were performed at the Children’s Hospital of Philadelphia using the Illumina genome-wide GSAMD-24 v2.0 SNP microarray. Approval for this study was obtained from the Stanford Institutional Review Board panel, and informed consent was obtained from all participants.

### Generation of hCO and hMHO

Cells were grown at ~80-90% confluency and detached using Accutase (StemCell Technologies, Cat. No. 07920) after 7 min of incubation at 37 °C. Accutase was quenched by adding E8 media (Thermo Fisher, A1517001) and spun down at 200 x g for 4 mins. The media was aspirated and cells were resuspended in 1-2 ml of E8 and counted using Trypan Blue (Thermo Fisher, T10282) staining in a Countess™ Automated Cell Counter (Invitrogen). Cells were transferred to a 15 ml conical tube and centrifuged at 200 x g for 4 mins before being resuspended in E8 with ROCK inhibitor (Y27, EMD Millipore, SCM075). 3×10^6^ cells were transferred to each well of a 24-well Aggrewell 800 plate (StemCell Technologies, 34815) and centrifuged at 100 x g for 3 mins.

Cortical organoids were generated as previously described^41^. Midbrain-hindbrain organoids were dislodged into Essential 6 media (E6) (Thermo Fisher, A1516401) with 2.5 *µ*m DM (dorsomorphin, Sigma-Aldrich, P5499) and 10 *µ*m SB (SB-431542, R&D Systems, 1614) on the day after aggregation. From day 3 to 5, E6 media was supplemented with 2.5 *µ*m DM, 10 *µ*m SB and 3 *µ*m CHIR (CHIR-99021, Selleck Chemicals, S1263). On day 6, media was changed to NPC media (Neurobasal media (Thermo Fisher Scientific, 10888022), B-27 supplement minus vitamin A (Thermo Fisher Scientific, 12587010), GlutaMAX (1:100; Thermo Fisher Scientific, 35050079), and penicillin–streptomycin (10,000 U/ml, 1:100; Thermo Fisher Scientific, 15140122) and supplemented with 1.5 *µ*m DM, 2.5 *µ*m SB, 3 *µ*m CHIR and 100 nM Smoothened agonist (SAG, EMD Millipore, 566660). On day 10, media was switched to NPC supplemented with 10 ng/ml Fibroblast growth factor 4 (FGF4, Peprotech, 100-31), 1.5 *µ*m CHIR and 100 nM SAG with daily media changes until day 15, and media changes every other day until day 23. On day 25, NPC media was supplemented 10 ng/ml Insulin-like growth factor 1 (IGF-1, Preprotech, 100-11), 20 ng/ml neurotrophin-3 (NT-3, PeproTech, 450-03), 20 ng/ml brain derived neurotrophic factor (BDNF, PeproTech, 450-02), 100 *µ*m cAMP (MilliporeSigma, D0627), 10 *µ*m DHA (MilliporeSigma, D2534), 200 *µ*m ascorbic acid (AA, Wako, 323-44822) and 2.5 *µ*m DAPT (StemCell Technologies, 72082) with media changes every other day until day 29. From day 31, the same media was used but without the addition of DAPT. From day 37, media was changed to NPC with B27 Plus Supplement (Thermo Fisher Scientific, A3582801) with the same factors used. From, day 47 media was changed to NPC with B27 Plus Supplement and media was changed every 4 days.

### RT-qPCR

Multiple organoids (3-5) were collected into one tube and considered as one sample. RNA was isolated using the RNeasy Plus Mini kit (Qiagen, 74136) and cDNA was synthesized using SuperScript^TM^ III First-Strand Synthesis Super Mix for qRT-PCR (Thermo Fisher Scientific, 11752250). qPCR was conducted using SYBR Green PCR Master Mix (Thermo Fisher Scientific, 4212704) using the QuantStudio^TM^ 6 Flex Real-Time PCR System (Thermo Fisher Scientific, 4485689). Primer sequences are provided in Supplementary table 1.

### Cryopreservation and immunocytochemistry

Organoids were fixed in 4% paraformaldehyde (PFA, Electron Microscopy Sciences, 15710) in DPBS overnight and subsequently washed three times in DPBS the next day. Fixed organoids were transferred to a solution containing 30% sucrose and left overnight at 4ºC, or until the organoids sank in solution. Samples were then transferred to a 1:1 mixture of OCT (Tissue-Tek OCT Compound 4583, Sakura Finetek) and 30% sucrose, and blocks were frozen using dry ice. 20 *µ*m thick sections were sectioned onto Super-frost^®^ Plus microscope slides (VWR, 48311-703) using a cryostat (Leica). For immunocytochemistry, cryosections were then blocked for 1 hr at room temperature (1% BSA (Sigma Aldrich, A9418-50G), 0.1% Triton X-100 (Thermo Scientific, AC327372500), 0.02% sodium azide (Sigma Aldrich, RTC000068-1L) diluted in DPBS). Samples were subsequently incubated overnight at 4°C with primary antibodies in blocking solution. The next day, cryosections were washed with blocking solution and then incubated with Alexa Fluor secondary antibodies (1:1000 dilution in blocking solution, donkey anti-rabbit IgG (H+L) Alexa Fluor 568 (Thermo Fisher Scientific, A-10042)) for 1 hr at room temperature. After incubation with secondary antibodies, sections were washed twice with DPBS and nuclei were visualized with Hoechst 33258 (Life Technologies, H3569). Sections were washed once more with DPBS and cover glass was mounted overnight using Aqua-Poly/Mount (Polysciences, 18606). The following antibodies were used for immunostaining: anti-TPH2 antibody (1:500 dilution, rabbit, Novus Biologicals, NB100-74555), anti-5-HT antibody (1:1000 dilution, rabbit, Immunostar, 20080), and anti-GFP antibody (1:1000 dilution, rabbit, Invitrogen, A-21311). Immunostained sections were imaged on an inverted confocal microscope (Leica Stellaris 5). Images were processed and analyzed using Fiji (ver. 2.14.0/1.54f).

### Preparation of scRNA-seq libraries and data analysis

Eight organoids were pooled for dissociation and dissociated using a Papain-based method as previously described^39^. Briefly, organoids were washed in PBS, chopped using a razor blade and incubated in a 6-well plate in Papain solution (30 U ml−1 Papain (Worthington Biochemical, LS003127), 1x Earle’s balanced salt solution (Sigma-Aldrich, E7510), 0.46% D(+)-glucose, 0.5 mM ethylenediaminetetraacetic acid (EDTA), 26 mM NaHCO_3_, 125 U ml−1 Deoxyribonuclease I (Worthington Biochemical, LS002007) and 6.1 mM L-cysteine (Sigma-Aldrich, C7880)) for 45 mins at 37°C with swirling in between. Then, organoids were transferred to a new 15 ml conical tube and the enzyme solution was removed and replaced by an enzyme inhibitor solution (2% trypsin inhibitor, Worthington Biochemical, LS00308, 1x Earle’s balanced salt solution, 0.46% D(+)-glucose, 26 mM NaHCO_3_, 125 U ml−1 Deoxyribonuclease I) and spun down at 200 x g for 5 mins. Organoids were subjected to multiple rounds of trituration and centrifugation with P1000 and P200 pipette tips until they were fully dissociated. Cells were resuspended in PBS + 0.1% BSA and filtered through a Flowmi^®^ pipette tip filter (Bel-Art, H13680-0040). Cell number and viability was assessed using a Countess Automated Cell counter with Trypan Blue staining.

Single-cell experiments were performed using the 10x Genomics Chromium platform using the 3’ end preparation protocol (10x Genomics, 1000123). Briefly, 7000 cells were targeted for each sample when loading cells on the microfluidic chip to generate GEMs. cDNA quality and quantity were assessed using the Agilent Bioanalyzer High Sensitivity DNA Assay. Single index primers were used for library prep (10x Genomics, 1000190, 10x Genomics, 1000213).

Sequencing libraries were sequenced on an Illumina NovaSeq S4 platform in 2×150 bp paired end mode at Admera Health. Demultiplexed fastq files were provided by the sequencing service for further processing. Raw fastq files were processed using 10x CellRanger Cloud services (v.5.0.0.) mapping to the GRCh2020-A human reference genome.

Processed data matrices were analyzed using Seurat^18^ (v4.1.1, v5.3.0) in R (v4.0.2, v4.5.0). Libraries were merged using Seurat’s merge function and quality was assessed. Cells expressing less than 500 or more than 7000 genes, more than 20,000 UMIs, or fraction of mitochondrial reads higher than 15% were removed from further analysis. The merged object was normalized using the LogNormalization() method and variable features were obtained using the VariableFeatures() function with the vst method with 3000 genes. Data were scaled using the ScaleData() function. The first 20 PCs were used for the FindNeighbors() function and FindClusters() with 0.5 resolution parameter. Cells types were annotated based on known marker gene expression and by using the FindMarkers() function in Seurat with min.pct = 0.3 and logfc.threshold = 0.3). Cells were annotated based on the following markers and similar clusters were merged for cell type annotation: hindbrain neural progenitors for *SOX2*^*+*^, *GATA2*^*+*,^ *GATA3*^*+*^, *NKX6-1*^*+*^, *NKX2-1*^*+*^, early neurons (*STMN2*^*+*^), glutamatergic neurons (*SCL17A6*^*+*^, *SCL17A7*^*+*^, *SLC17A8*^*+*^), GABAergic neurons (*GAD1*^*+*^, *GAD2*^*+*^), dopaminergic neurons (*TH+*), serotonergic neurons (*TPH2+, SLC6A4*^*+*^, *FEV*^*+*^), cycling cells (*MKI67*^*+*^) and astroglia (*SOX9*^*+*^, *AQP4*^*+*^). One cluster positive for neuronal marker *STMN2*^*+*^*and PHOX2B*^*+*^ and less distinct neurotransmitter expression was labeled as ‘Other neurons’. A small cluster of mesenchymal-like cells containing 72 cells expressing *COL1A1*^*+*^ *and LUM*^*+*^ were removed from the UMAP embedding.

A downsampled data set of the human developing brain transcriptome^8,16^ was used to annotate the hMHO data set using the Seurat label transfer function. First, the label transfer was performed on the whole reference data set to obtain the best matching developmental time point in the reference to perform further mapping. This was achieved by using the FindingVariableFeatures() function using the ‘vst’ method and the top 3000 features for both datasets, followed by computing anchors between the reference primary tissue data set and query organoid data using the FindTransferAnchors() function using the top 30 PCs. Predictions were calculated using TransferData() on the ‘Dataset’ variable in the reference dataset and using the top 30 PCs. Then, data from the primary tissue reference were subset to PCW 6.9 based on highest prediction score mapping of the cells to the hMHO data. Label transfer was then performed to obtain information about the brain regional mapping. Predictions were calculated based on the ‘Subregion’ variable in the primary tissue reference data set using 30 PCs and cells were labeled according to the maximum prediction score. Labels “Thalamus”, “Striatum” and “Cortex” were summarized into the label “Forebrain”.

The 5-HT neuron cluster was subset and reclustered for further analysis. Briefly, data were normalized using NormalizeData() using log normalization, variable features were identified using the ‘vst’ method in FindVariableFeatures with 3000 features and data were scaled using ScaleData(). The top 20 PCs were used for FindNeighbors() and FindClusters() was run with a resolution of 0.5. Markers for further annotation of the data were obtained from Okaty *et al*.^19^. Cells were further annotated to identify brain regional mapping using label transfer as described above. Briefly, the cells from the 5-HT subcluster were used as query and the 6.9 PCW primary tissue dataset as reference using 30 PCs for running TransferData() and calculating predictions based on the ‘Subregion’ variable. The cells were labeled based on the maximum prediction score. Cells were mapped to an E13 mouse reference spatial atlas using VoxHunt^20^ by using the top 10 markers for each region of the refence data set and similarity correlation of cells was plotted in sagittal view on the mouse spatial reference.

### Viral labeling

AAV-SYN1::mCherry (AAV-DJ-hSyn1-mCherry, GVVC-AAV-017) and AAV-EF1α::DIO-ChR2-eYFP (AAV-DJ-EF1-DIO hChR2(H134R)-eYFP, GVVC-AAV-038) were obtained from the Stanford University Gene Vector Virus Core. The AAV-TPH2::EGFP-WPRE and AAV-TPH2::Cre were designed based on the promoter sequence published in Gentile *et al*.^26^ and were obtained from VectorBuilder. LV-SYN1::PsychLight2 was based on Dong *et al*.^26^ and obtained from VectorBuilder. AAV-SYN1-GCaMP7s was obtained from Addgene (104487-AAV1).

Viral infections were conducted as previously described^42^. Briefly, 3-4 organoids were infected with AAV or lentivirus in 200 µl of NPC with B27 Plus in one well of a 24-well plate in a 1:1000 dilution of virus. Plates were left tilted in the incubator overnight, and 800 µl of fresh media was added the next day. Infected organoids were washed twice with 1.5 ml fresh NPC media with B27 Plus on the following day. Alternatively, neural organoids were transferred into a 1.5 mL Eppendorf tube containing 200 µl of neural media and incubated with the virus overnight at 37°C, 5% CO2 before washing steps. The following day, neural organoids were transferred into fresh culture media in ultra-low attachment plates (Corning, 3471, 3261).

### Generation of hNMA

To generate neuromodulatory assembloids, hMHO and hCO were generated individually from hiPS cells as described above. After day 50, hMHO and hCO were assembled by placing them in close proximity in 1.5 ml Eppendorf tubes, inside an incubator, for 3 days or by placing them in tilted 24-well ultra-low attachment plates (3437, Corning). Medium was gently replaced on day 2. On the third day, assembloids were transferred using a cut P1000 pipette tip to 24-well ultra-low attachment plates in the neural media described above. Media was changed every 4 days.

### Projection imaging

TPH2::EYFP^+^ cells projecting into hCO from hMHO and SYN1::mCherry^+^ cells projecting to hMHO from hCO were imaged under environmentally controlled conditions in hNMA using a Leica TCS SP8 or Leica Stellaris 5 confocal microscope with a motorized stage. Assembloids were transferred to a well in a 24-well glass bottom plate (Cellvis, P24-0-N) in culture medium, and images were taken using a 10x objective to capture the entire hNMA at a depth of 100 *µ*m (22q11.2DS imaging). Projections were quantified using Fiji (ImageJ, version 2.1.0, NIH). ROIs were manually drawn to cover the area of the hMHO or hCO side in maximal projection confocal stacks. The percentage of fluorescence positive pixels over total area of hMHO or hCO was calculated in binary images with consistent threshold values across images. All images related to 22q11.2DS experiments were blinded during image analysis.

### Serotonin sensor imaging

For live cell imaging, labeled hCO, hMHO or hNMA were transferred into a well of a Corning™ 96-Well microplate (Corning, 4580) in 150 µl of neural media or on a 20 mm glass coverslip in a custom made perfusion chamber and incubated in an environmentally controlled chamber for 15–30 min before imaging on a Leica TCS SP8 or Leica Stellaris 5 confocal microscope.

Dose-response experiments were performed using a peristaltic perfusion system with neural organoids in the perfusion chamber (PPS2, Multi Channel Systems MCS). Organoids were perfused with 1 ml of each dosage of 5-HT (1 nM to 100 *µ*m, Sigma Aldrich, H9523) followed by 100 *µ*m EMD 281014 (Tocris, 4470). Drug addition experiments were performed by adding the final 30 mM KCl with or without 100 *µ*m fluoxetine (Tocris, 0925) directly into the imaging well on 96-well microplates. Both imaging sessions were conducted using 465 nm laser and a 20x air objective on a Leica microscope (SP8 and Leica Stellaris 5). PsychLight2 was imaged at 0.3 frames per second in timelapse mode (xyt). Results were analyzed with Fiji (NIH, v2.14.0/1.54f). After ROI registration, the raw time series were transformed into relative changes in fluorescence: ΔF/F(t) = (F(t) − F_0_)/F_0_ for quantifications. All serotonin sensor imaging experiments related to 22q11.2DS experiments were blinded during image acquisition and image analysis.

### In vitro whole cell recordings

Midbrain-hindbrain organoids were infected with an AAV-TPH2::EGFP to label 5-HT neurons, and hCO were infected with an AAV-SYN1::mCherry to label cortical neurons. For assembloid recordings, hCO were infected with AAV-SYN1::mCherry and hMHO with AAV-TPH2::Cre + AAV-EF1α::DIO-ChR2-EYFP around day 60 and were fused around day 70 to generate hNMA. Assembloids were typically fused for around 60 days before performing whole cell patch recording on SYN1::mCherry^+^ neurons.

Patch-clamp recordings were conducted, as described previously^30,39^. Briefly, assembloids or organoids were placed onto cell culture inserts (0.4-*µ*m pore size, Corning, 353090) positioned in six-well plates to form flattened assembloids. GFP^+^ neurons in hMHO or mCherry^+^ neurons in hCO were identified and patched. Whole-cell recordings were performed at room temperature and were visualized using an upright microscope (Scientifica) equipped with an INFINITY2 charge-coupled device camera and INFINITY Capture Software (Teledyne Lumenera). The external aCSF recording solution contained 124 mM NaCl, 3 mM KCl, 1.25 mM NaH_2_PO_4_, 1 mM MgSO_4_, 2 mM CaCl_2_, 26 mM NaHCO_3_ and 10 mM D-(+)-glucose. During recordings, organoids and assembloids were perfused with aCSF recording solution (bubbled with 95% O_2_ and 5% CO_2_). Thick-walled borosilicate pipettes (resistance of 5–9 MΩ) were filled with an internal solution consisting of 135 mM K-gluconate, 20 mM KCl, 0.1 mM EGTA, 2 mM MgCl_2_, 2 mM sodium-ATP, 10 mM HEPES and 0.3 mM sodium-GTP (pH adjusted to 7.28 with KOH; approximately 302 mOsm). Data were acquired using a MultiClamp 700B Amplifier (Molecular Devices, Clampex 10.7) and a Digidata 1550B Digitizer (Molecular Devices). Data were low-pass filtered at 2 kHz and digitized at 20 kHz.

Spontaneous excitatory postsynaptic currents (sEPSCs) were recorded at command voltage of –60 mV in the voltage-clamp mode before liquid junction potential (LJP) correction. Evoked action potentials (eAPs) were recorded at above the action potential threshold potential for each patched neuron (around −54 mV) in the current clamp mode. For action potential *F*–*I* curves, cells were current-clamped at −60 mV and current steps (1 s duration) were given with an increment of 20 pA with a 10 s interval. The LJP was calculated using JPCalc and recording data (resting membrane potential and action potential threshold voltage) were corrected with an estimated of ~14-mV LJP offline.

To study the effect of 5-HT on evoked APs in hMHO, 100 *µ*m 5-HT (Sigma Aldrich, H9523, freshly prepared on day of recording) or 50 *µ*m 5-HT1A receptor antagonist NAN-190 (Tocris, 0553) were applied via bath solution perfusion. Evoked action potential frequency was measured as the average of 100 s before addition of 5-HT. 5-HT was added and 60 s after addition, the response was quantified as the average frequency over 100 s. The same quantification intervals were used for NAN-190 addition.

To study the functional connection in the hNMA, light train stimulations at 5ms/20Hz were delivered by whole-field light-emitting diode illumination (460 nm; maximal power ~20mW; CoolLED) and applied through a 40x objective. Average baseline frequency of sEPSC was measured as the average over at least 90 s prior to light stimulation. One min after light stimulation, sEPSC frequency was averaged over a time interval of 2 mins. Slow inward current responses were defined as a change of at least 5 pA from baseline within 10 mins after light stimulation. All recording data were further analyzed by Clampfit (v.10.7; Molecular Devices) and eFEL python package (v. 5.7.16)^43^.

### Biocytin labeling and imaging

TPH2-EGFP^+^ cells in hMHO were labeled during electrophysiological measurements by adding 0.2% Biocytin (Sigma Aldrich, B4261) to the internal solution and injecting into cells. Organoids were subsequently fixed in 4% PFA at 4°C over night and washed three times with DPBS the following day. Samples were then stained in blocking solution (DPBS + 10% normal donkey serum (Sigma Aldrich, S30-M) + 0.3% TritonX) with 1:1000 Streptavidin DyLight549 (Vector Labs, SA05549) and incubated over night at 4°C. Samples were washed with DPBS the following day, embedded between two cover slips in Aqua/PolyMount mounting media (18606, Polysciences) and imaged on a Leica SP8 at 10x magnification using z-stacks. Maximum projection of z-stack images was performed in Fiji, thresholded and binarized to visualize the morphology of the 5-HT neurons.

### Live calcium imaging in hCO and hNMA

Cortical organoids were infected with AAV-SYN1::GCaMP7s (Addgene, 104487-AAV1) around day 55 and fused to hMHO at approximately day 60. Calcium imaging was performed one month after fusion of organoids. On the day of imaging, assembloids were transferred to fresh Neurobasal media with B27 Plus and allowed to equilibrate in a 37°C incubator with 5% CO2 for 1 hour before imaging. The assembloids were then transferred to an inverted confocal microscope (Leica SP8) with temperature control set at 37°C. Imaging was performed using the 10x objective and the resonant scanner with a line frequency of 8 kHz and a frame resolution of 1024 x 1024 pixels. Imaging was performed for 5 minutes at a scan zoom of 1.25x with bidirectional scanning. Time-lapse images were processed and analyzed using Fiji (ver. 2.14.0/1.54f). Images were processed using the Gaussian Blur 3D function with the following parameters: X sigma = 0.0, Y sigma = 0.0, Z sigma = 2.0. Cell bodies were selected using the elliptical tool and added to the ROI manager. Once all active cells had been selected, fluorescence intensity over time was measured using the Multi Measure tool with “Mean gray value” selected. The intensity values for each cell were used as input for scaled correlation analysis (SCA)^29,30^ to measure maximum peak correlation across cells in organoids. Only organoids with at least 3 active cells were considered for the analysis.

### HPLC measurements

3-5 organoids were collected into a 1.5 ml Eppendorf tube for HPLC sample preparation. All leftover media was removed and 100 µl of 0.2 M perchlorid acid was added to the organoids before homogenization using a pellet pestle (DWK Life Sciences, 749521-1500). Samples were incubated on ice for 30 min and centrifuged for 15 mins at 21,100 x g at 4 °C. The supernatant was filtered using Minisart filters (Sartorius, 60-097), frozen on dry ice and stored at −80°C until further use. Tissue pellets were frozen on dry ice and stored at −80°C.

5-HT levels in the samples were determined by HPLC with electrochemical detection. Briefly, the supernatant was injected into the column and the HPLC mobile phase (0.1 M ammonium acetate buffer, 50 mg/l EDTA, 0.05 M sodium sulfate, 30% MeOH, pH 6.0) was pumped through a chromatography column (CAX, Eicom) and a 20 µl sample loop at a flow rate of 0.25 ml/min. A graphite electrode (WE-3G, Eicom) set at a potential of 450 mV was used for electrochemical detection. Quantification was performed using a PowerChrom analysis system (ADInstruments) using external standards (Sigma) and the concentration was calculated by comparing the HPLC peak of samples with the peak area of known concentrations of the standards analyzed on the same day. The levels of 5-HT were normalized to the amount of total protein in the samples which was measured from the sample pellets using Pierce BCA assay (Thermo Fisher Scientific, 23227).

### Statistics

Values are given as mean ± standard error of the mean (SEM) unless otherwise stated. Comparisons between experimental groups were analyzed using two-tailed *t*-tests or their non-parametric equivalents. For more complex designs, one-way or two-way ANOVA tests or their non-parametric equivalents were used with multiple testing correction. *P* values of less than 0.05 were considered statistically significant. GraphPad Prism (10.5.0) was used for statistical analysis.

**Figure S1:**
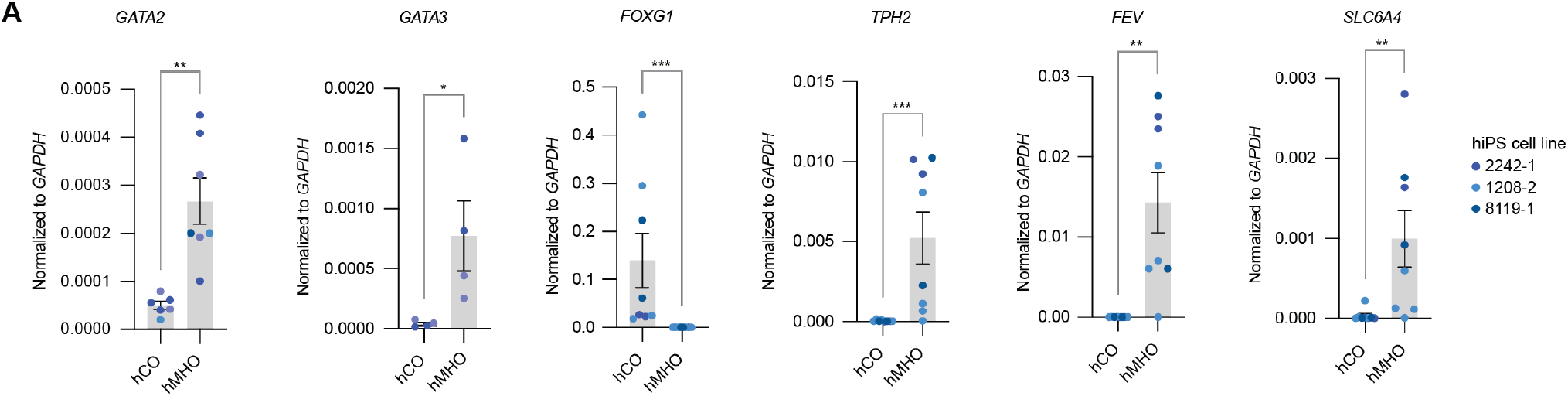
Marker gene expression in hMHO. **(A)** RT-qPCR of marker gene expression comparing hCO and hMHO. Day 19-22: *GATA2* (**p = 0.0012), *GATA3* (*p = 0.0286); day 50-60: *FOXG1* (***p = 0.0002), *TPH2* (***p = 0.0006), *FEV* (**p = 0.0047), *SLC6A4* (**p = 0.0070); (Mann-Whitney test for all genes, 3 hiPS cell lines, 1-4 differentiation experiments, Mann-Whitney test for all genes).

**Figure S2:**
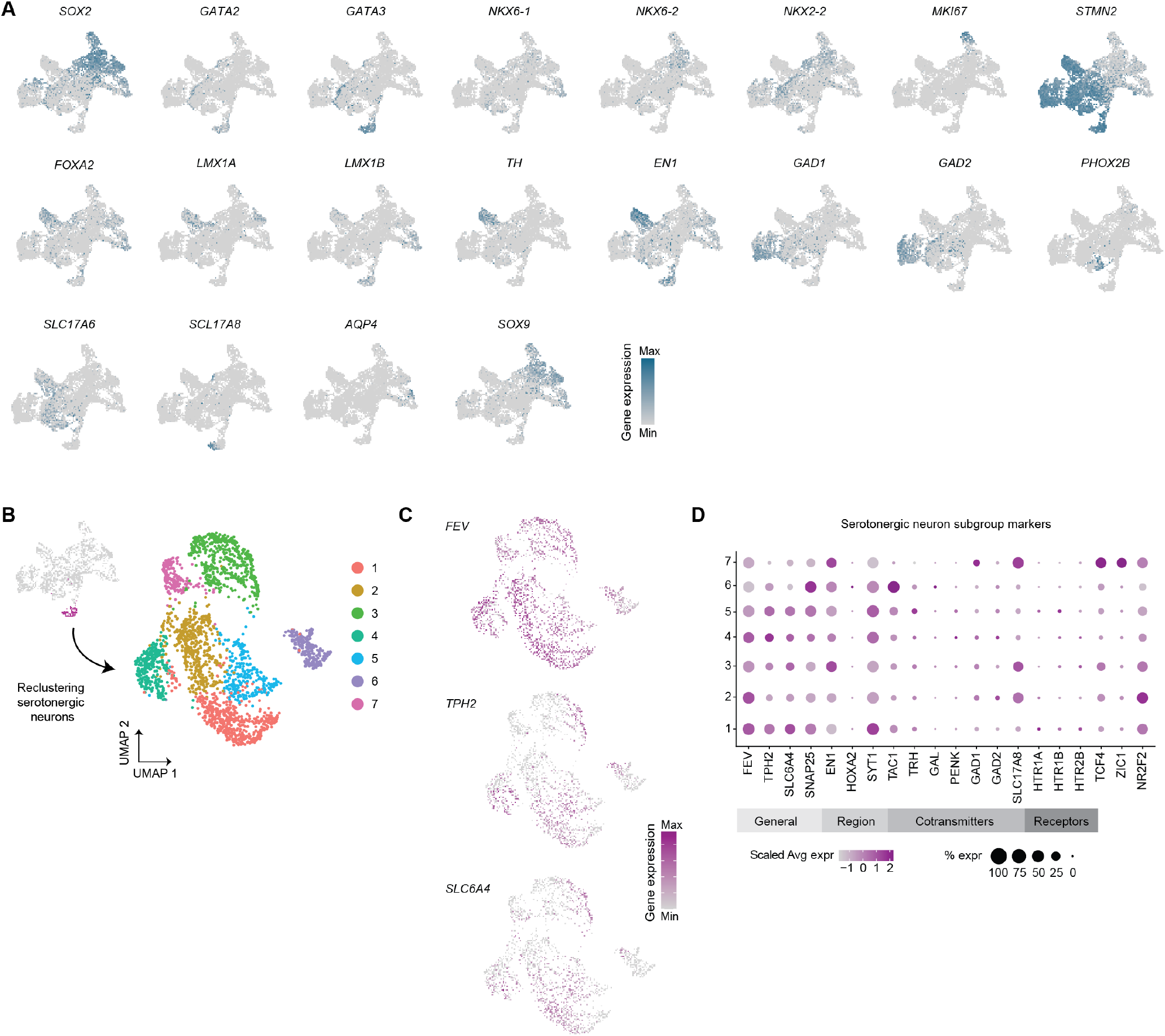
scRNA-seq characterization of hMHO. **(A)** Marker gene expression in hMHO. **(B)** Subclustering of 5-HT neurons. **(C)** Expression of canonical markers in 5-HT neurons. **(D)** Markers of 5-HT neuron subtypes.

**Figure S3:**
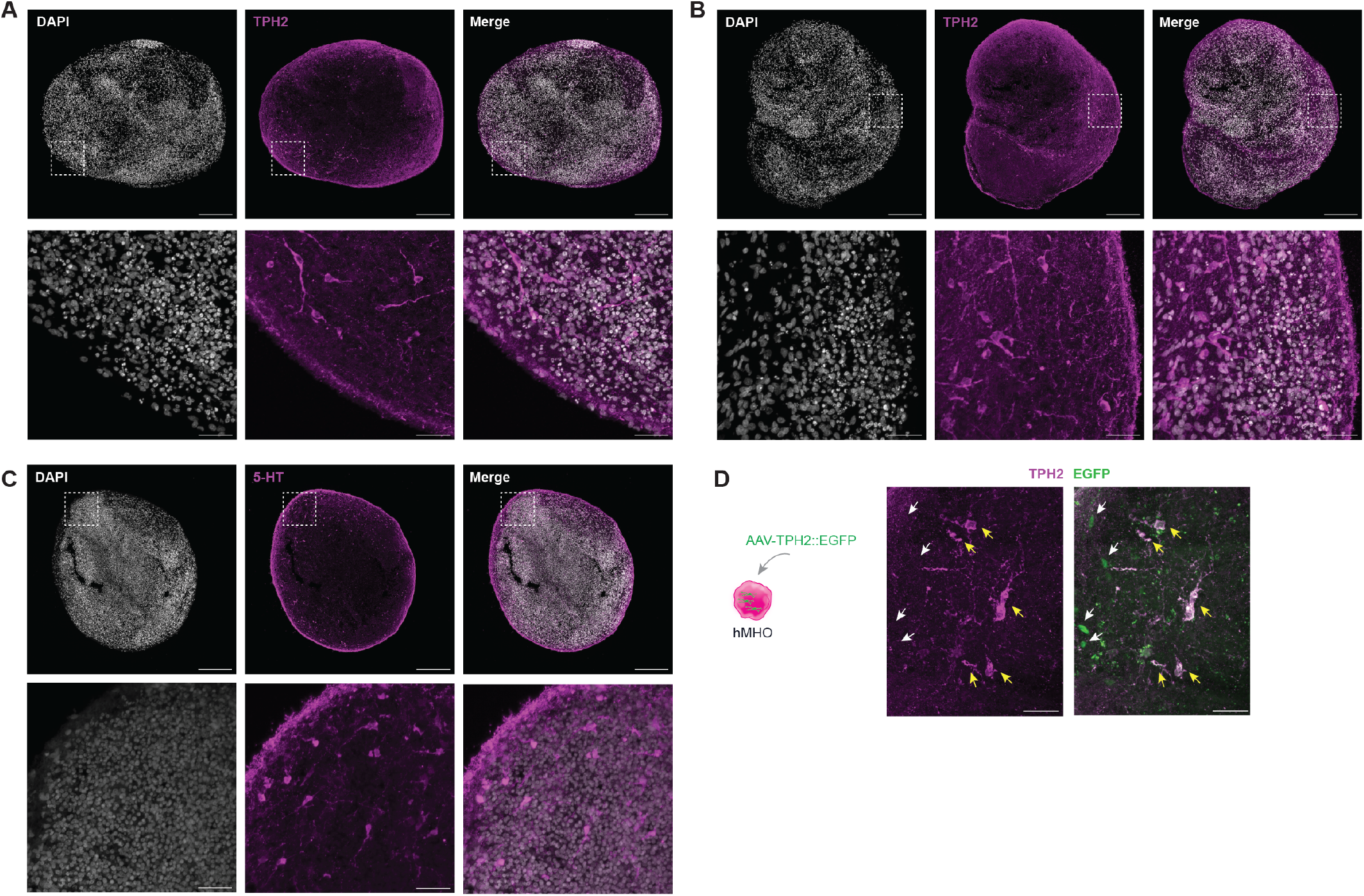
Immunohistochemical stainings of hMHO. **(A, B)** Representative immunohistochemical stainings of TPH2 in hMHO (top scale bar = 250 *µ*m; bottom scale bar = 40 *µ*m). **(C)** Representative immunohistochemical staining of 5-HT in hMHO (top scale bar = 250 *µ*m; bottom scale bar = 40 *µ*m). **(D)** IHC staining for TPH2 and EGFP in hMHO (d197) infected with AAV-TPH2::EGFP. Yellow arrows: EGFP^+^/TPH2^+^ cells, white arrows: EGFP^+^/TPH2^-^ cells (scale bar = 40 *µ*m).

**Figure S4:**
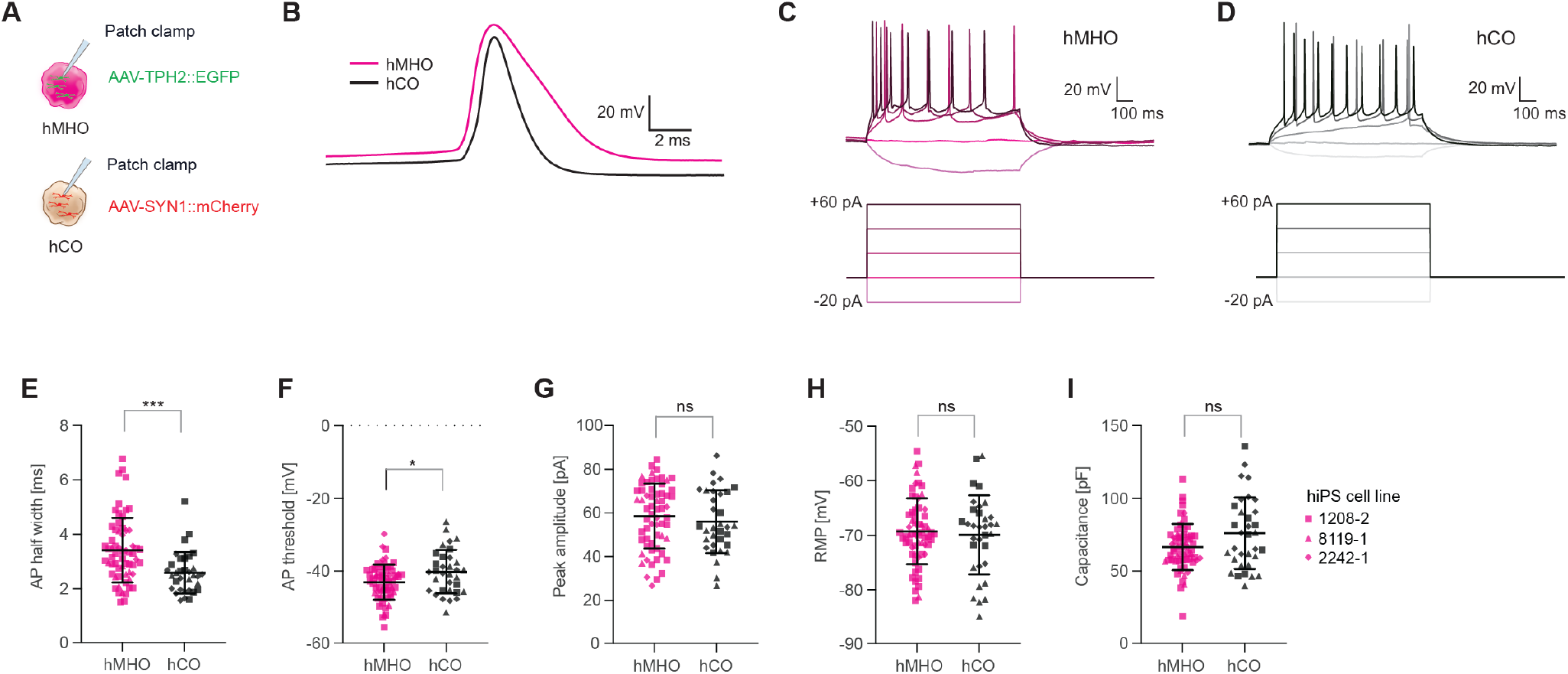
Electrophysiological characterization of hMHO and hCO. **(A)** Schematic of patch clamp recordings in TPH2::EGFP^+^ neurons in hMHO and SYN1::mCherry^+^ neurons in hCO. **(B)** Representative traces of action potentials in hMHO and hCO. **(C)** Example action potential traces in hMHO during current injection. **(D)** Example action potential traces in hCO during current injection. **(E)** Action potential half width of labeled cells in hMHO and hCO (hMHO: 62 cells, 6 differentiation experiments, 3 hiPS cell lines, day 133 - 205; hCO: 33 cells, 3 differentiation experiments, 3 hiPS cell lines, day 152 - 208. Mann Whitney test, ***p = 0.0002). **(F)** Action potential threshold of labeled cells in hMHO and hCO (hMHO: 62 cells, 6 differentiation experiments, 3 hiPS cell lines day 133 - 205; hCO: 33 cells, 3 differentiation experiments, 3 hiPS cell lines, day 152 - 208. Mann Whitney test, *p = 0.0355). **(G)** Peak action potential amplitude of labeled cells in hMHO and hCO (hMHO: 62 cells, 6 differentiation experiments, 3 hiPS cell lines, day 133 - 205; hCO: 33 cells, 3 differentiation experiments, 3 hiPS cell lines, day 152 - 208. Mann Whitney test, p = 0.3802). **(H)** Resting membrane potential of labeled cells in hMHO and hCO (hMHO: 62 cells, 6 differentiation experiments, 3 hiPS cell lines, day 133 - 205; hCO: 33 cells, 3 differentiation experiments, 3 hiPS cell lines, day 152 - 208. Mann Whitney test, p = 0.9829). **(I)** Capacitance of labeled cells in hMHO and hCO (hMHO: 62 cells, 6 differentiation experiments, 3 hiPS cell lines, day 133 - 205; hCO: 33 cells, 3 differentiation experiments, 3 hiPS cell lines, day 152 - 208. Mann Whitney test, p = 0.1197). All error bars are shown as mean ± SD.

**Figure S5:**
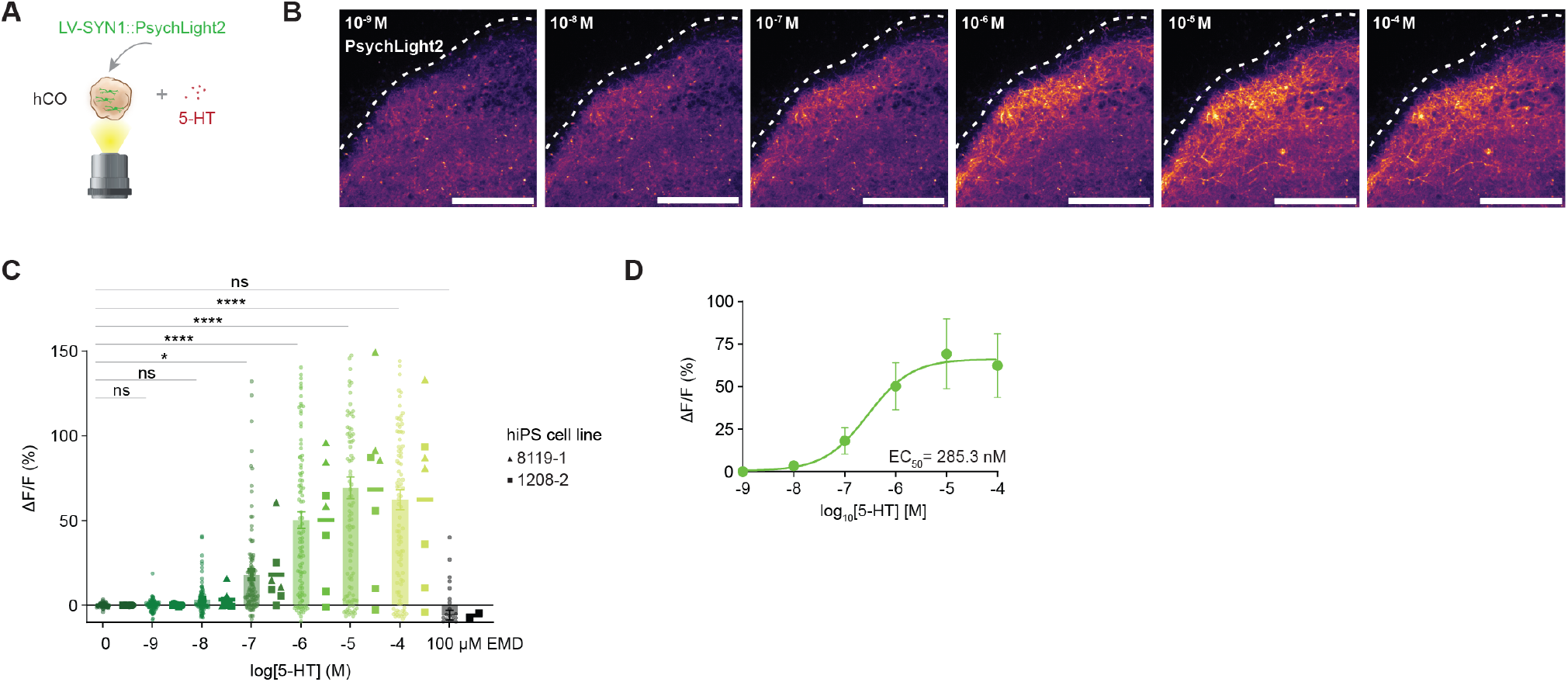
Serotonin sensor characterization in hCO. **(A)** Schematic of PsychLight2 sensor imaging in hCO with 5-HT addition. **(B)** Example images of PsychLight2 in hCO at increasing 5-HT concentrations. **(C)** Fluorescence intensity change during 5-HT and blocker addition across hiPS cell lines (30 - 105 ROIs (left), 2 - 7 hCO (right), 2 hiPS cell lines, 2 differentiation experiments. One-way ANOVA, multiple comparison testing compared to 0, *p = 0.0126, ****p<0.0001). **(D)** Dose-response curve of PsychLight2 fluorescence intensity in hCO during increasing concentrations of 5-HT (derived from panel C; 7 hCO, 2 hiPS cell lines, 2 differentiation experiments).

**Figure S6:**
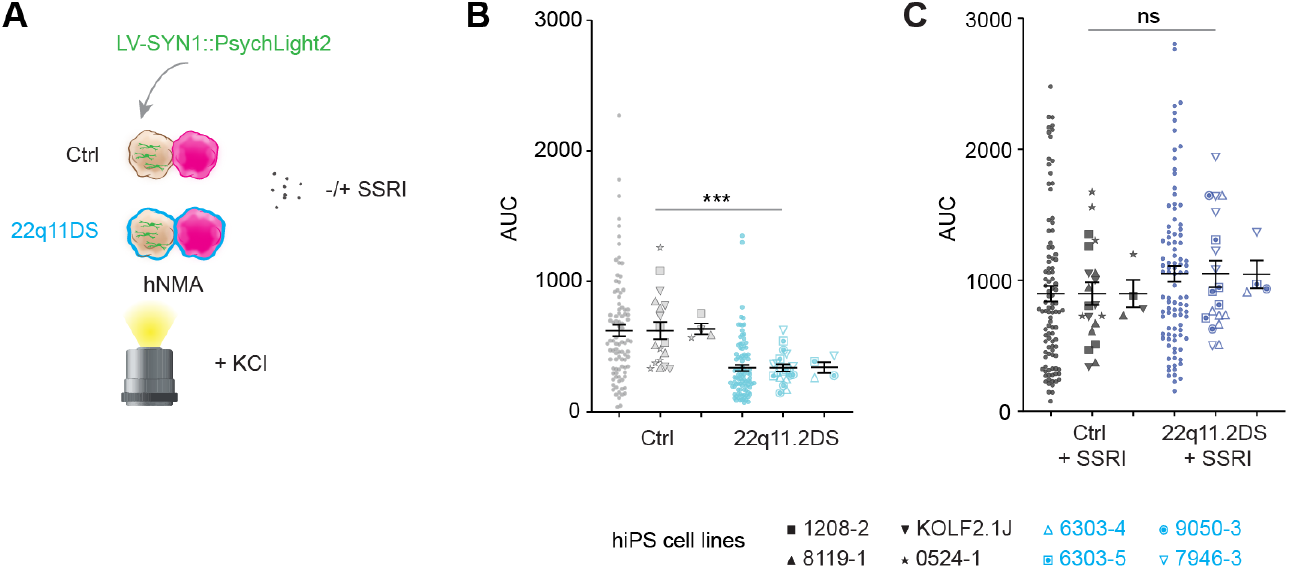
Serotonin sensor dynamics in 22q11.2DS. **(A)** Schematic of PsychLight2 sensor imaging in control and 22q11.2DS hCO with and without SSRI addition. **(B)** Quantification of AUC of PsychLight2 signal in hCO from control and 22q11.2DS in hNMA (Ctrl: 90 ROIs (left), 18 assembloids (middle), 4 hiPS cell lines (right), 3 differentiation experiments; 22q11.2DS: 95 ROIs (left), 19 assembloids (middle), 4 hiPS cell lines (right) 3 differentiation experiments. Unpaired t-test with Welch’s test, ***p=0.0006). **(C)** Quantification of AUC of PsychLight2 signal in hCO from control and 22q11.2DS in hNMA with SSRI addition (Ctrl: 100 ROIs (left), 20 assembloids (middle), 4 hiPS cell lines (right), 3 differentiation experiments; 22q11.2DS: 95 ROIs (left), 19 assembloids (middle), 4 hiPS cell lines (right), 3 differentiation experiments. Unpaired t-test with Welch’s test, p = 0.2638).

## References

1. Bonnin, A. et al. A transient placental source of serotonin for the fetal forebrain. Nature 472, 347–350 (2011).

2. Deneris, E. & Gaspar, P. Serotonin neuron development: shaping molecular and structural identities. Wiley Interdiscip. Rev. Dev. Biol. 7, (2018).

3. Takahashi, H., Nakashima, S., Ohama, E., Takeda, S. & Ikuta, F. Distribution of serotonin-containing cell bodies in the brainstem of the human fetus determined with immunohistochemistry using antiserotonin serum. Brain Dev. 8, 355–365 (1986).

4. Lesch, K.-P. & Waider, J. Serotonin in the Modulation of Neural Plasticity and Networks: Implications for Neurodevelopmental Disorders. Neuron 76, 175–191 (2012).

5. Celada, P., Puig, M. V. & Artigas, F. Serotonin modulation of cortical neurons and networks. Front. Integr. Neurosci. 7, 25 (2013).

6. Terry, A. V., Buccafusco, J. J. & Wilson, C. Cognitive dysfunction in neuropsychiatric disorders: selected serotonin receptor subtypes as therapeutic targets. Behav. Brain Res. 195, 30–38 (2008).

7. Hodge, R. D. et al. Conserved cell types with divergent features in human versus mouse cortex. Nature 573, 61–68 (2019).

8. Amin, N. D. et al. Generating human neural diversity with a multiplexed morphogen screen in organoids. Cell Stem Cell 31, 1831-1846.e9 (2024).

9. Kim, J.-I. et al. Human assembloids reveal the consequences of CACNA1G gene variants in the thalamocortical pathway. Neuron 112, 4048-4059.e7 (2024).

10. Birey, F. et al. Assembly of functionally integrated human forebrain spheroids. Nature 545, 54–59 (2017).

11. McDonald-McGinn, D. M. et al. 22q11.2 deletion syndrome. Nat. Rev. Dis. Primer 1, 15071 (2015).

12. Vorstman, J. A. S. et al. The 22q11.2 deletion in children: high rate of autistic disorders and early onset of psychotic symptoms. J. Am. Acad. Child Adolesc. Psychiatry 45, 1104–1113 (2006).

13. Wurst, W. & Bally-Cuif, L. Neural plate patterning: upstream and downstream of the isthmic organizer. Nat. Rev. Neurosci. 2, 99–108 (2001).

14. Lu, J. et al. Generation of serotonin neurons from human pluripotent stem cells. Nat. Biotechnol. 34, 89–94 (2016).

15. Valiulahi, P. et al. Generation of caudal-type serotonin neurons and hindbrain-fate organoids from hPSCs. Stem Cell Rep. 16, 1938–1952 (2021).

16. Braun, E. et al. Comprehensive cell atlas of the first-trimester developing human brain. Science 382, eadf1226 (2023).

17. Liu, C. et al. Pet-1 is required across different stages of life to regulate serotonergic function. Nat. Neurosci. 13, 1190–1198 (2010).

18. Hao, Y. et al. Dictionary learning for integrative, multimodal and scalable single-cell analysis. Nat. Biotechnol. 42, 293–304 (2024).

19. Okaty, B. W., Commons, K. G. & Dymecki, S. M. Embracing diversity in the 5-HT neuronal system. Nat. Rev. Neurosci. 20, 397–424 (2019).

20. Fleck, J. S. et al. Resolving organoid brain region identities by mapping single-cell genomic data to reference atlases. Cell Stem Cell 28, 1148-1159.e8 (2021).

21. Gentile, M. T. et al. Tryptophan hydroxylase 2 (TPH2) in a neuronal cell line: modulation by cell differentiation and NRSF/rest activity. J. Neurochem. 123, 963–970 (2012).

22. Liu, R.-J., Lambe, E. K. & Aghajanian, G. K. Somatodendritic autoreceptor regulation of serotonergic neurons: dependence on L-tryptophan and tryptophan hydroxylase-activating kinases. Eur. J. Neurosci. 21, 945–958 (2005).

23. Aghajanian, G. K. & Lakoski, J. M. Hyperpolarization of serotonergic neurons by serotonin and LSD: studies in brain slices showing increased K+ conductance. Brain Res. 305, 181–185 (1984).

24. Kirby, L. G., Pernar, L., Valentino, R. J. & Beck, S. G. Distinguishing characteristics of serotonin and non-serotonin-containing cells in the dorsal raphe nucleus: electrophysiological and immunohistochemical studies. Neuroscience 116, 669–683 (2003).

25. Li, Y. Q., Li, H., Kaneko, T. & Mizuno, N. Morphological features and electrophysiological properties of serotonergic and non-serotonergic projection neurons in the dorsal raphe nucleus. An intracellular recording and labeling study in rat brain slices. Brain Res. 900, 110–118 (2001).

26. Dong, C. et al. Psychedelic-inspired drug discovery using an engineered biosensor. Cell 184, 2779-2792.e18 (2021).

27. Schmitz, N., Hodzic, S. & Riedemann, T. Common and contrasting effects of 5-HTergic signaling in pyramidal cells and SOM interneurons of the mouse cortex. Neuropsychopharmacol. Off. Publ. Am. Coll. Neuropsychopharmacol. 50, 783–797 (2025).

28. Sengupta, A., Bocchio, M., Bannerman, D. M., Sharp, T. & Capogna, M. Control of Amygdala Circuits by 5-HT Neurons via 5-HT and Glutamate Cotransmission. J. Neurosci. 37, 1785–1796 (2017).

29. Nikolić, D., Mureşan, R. C., Feng, W. & Singer, W. Scaled correlation analysis: a better way to compute a cross-correlogram. Eur. J. Neurosci. 35, 742–762 (2012).

30. Kim, J.-I. et al. Human assembloid model of the ascending neural sensory pathway. Nature 642, 143–153 (2025).

31. Evers, L. J. M. et al. Serotonergic, noradrenergic and dopaminergic markers are related to cognitive function in adults with 22q11 deletion syndrome. Int. J. Neuropsychopharmacol. 17, 1159–1165 (2014).

32. Khan, T. A. et al. Neuronal defects in a human cellular model of 22q11.2 deletion syndrome. Nat. Med. 26, 1888–1898 (2020).

33. Mancini, V. et al. Long-term effects of early treatment with SSRIs on cognition and brain development in individuals with 22q11.2 deletion syndrome. Transl. Psychiatry 11, 336 (2021).

34. Farrelly, L. A. et al. Histone serotonylation is a permissive modification that enhances TFIID binding to H3K4me3. Nature 567, 535–539 (2019).

35. Ciranna, L. Serotonin as a modulator of glutamate- and GABA-mediated neurotransmission: implications in physiological functions and in pathology. Curr. Neuropharmacol. 4, 101–114 (2006).

36. Shin, D. et al. Thalamocortical organoids enable in vitro modeling of 22q11.2 microdeletion associated with neuropsychiatric disorders. Cell Stem Cell 31, 421-432.e8 (2024).

37. Mosheva, M., Korotkin, L., Gur, R. E., Weizman, A. & Gothelf, D. Effectiveness and side effects of psychopharmacotherapy in individuals with 22q11.2 deletion syndrome with comorbid psychiatric disorders: a systematic review. Eur. Child Adolesc. Psychiatry 29, 1035–1048 (2020).

38. Gebauer, L., Jensen, O., Neif, M., Brockmöller, J. & Dücker, C. Overlap and Specificity in the Substrate Spectra of Human Monoamine Transporters and Organic Cation Transporters 1, 2, and 3. Int. J. Mol. Sci. 22, 12816 (2021).

39. Miura, Y. et al. Generation of human striatal organoids and cortico-striatal assembloids from human pluripotent stem cells. Nat. Biotechnol. 38, 1421–1430 (2020).

40. Pantazis, C. B. et al. A reference human induced pluripotent stem cell line for large-scale collaborative studies. Cell Stem Cell 29, 1685-1702.e22 (2022).

41. Yoon, S.-J. et al. Reliability of human cortical organoid generation. Nat. Methods 16, 75–78 (2019).

42. Miura, Y. et al. Engineering brain assembloids to interrogate human neural circuits. Nat. Protoc. 17, 15–35 (2022).

43. Ranjan, R. et al. eFEL. Zenodo 10.5281/ZENODO.593869 (2024).

